# Influence of Demographic, Socio-economic, and Brain Structural Factors on Adolescent Neurocognition: A Correlation Analysis in the ABCD Initiative

**DOI:** 10.1101/2023.02.24.529930

**Authors:** Mohammad Arafat Hussain, Grace Li, Ellen Grant, Yangming Ou

**Affiliations:** Department of Pediatrics, Boston Children’s Hospital, Harvard Medical School, 401 Park Drive, Boston, MA 02115; Harvard College, Cambridge, MA 02138; Computational Health Informatics Program, Boston Children’s Hospital, Harvard Medical School, 401 Park Drive, Boston, MA 02115; Department of Radiology, Harvard Medical School, 401 Park Drive, Boston, MA 02115

## Abstract

The Adolescent Brain Cognitive Development (ABCD) initiative is a longitudinal study aimed at characterizing brain development from childhood through adolescence and identifying key biological and environmental factors that influence this development. The study measures neurocognitive abilities across a multidimensional array of functions, with a focus on the critical period of adolescence during which physical and socio-emotional changes occur and the structure of the cortical and white matter changes. In this study, we perform a correlation analysis to examine the linear relation of adolescent neurocognition functions with the demographic, socio-economic, and magnetic resonance imaging-based brain structural factors. The overall goal is to obtain a comprehensive understanding of how natural and nurtural factors influence adolescent neurocognition. Our results on *>* 10,000 adolescents show many positive and negative statistical significance interrelations of different neurocognitive functions with the demographic, socioeconomic, and brain structural factors, and also open up questions inviting further future studies.

## Introduction

The Adolescent Brain Cognitive Development (ABCD) initiative^1^ recruited children 9-10 years of age to perform a longitudinal study. This study aims to characterize brain development from childhood through adolescence and to find the key biological and environmental factors that influence this development. ABCD consortium aimed to measure neurocognitive abilities across a multidimensional array of functions^1^ as summarized in Table 1^2^. The variation in general acuity of neurocognition in humans is the result of the efficiency of the subdomain (listed in the fourth column of Table 1)^3^. Adolescence is a critical period of time in which physical and socio-emotional changes occur in humans, which greatly shape the level of acuity in neurocognitive subdomains^2^. More importantly, this period involves changes in the structure of the cortical and white matter. However, the onset of many mental illnesses is also recorded in this period of life^4^.

**Table 1.**
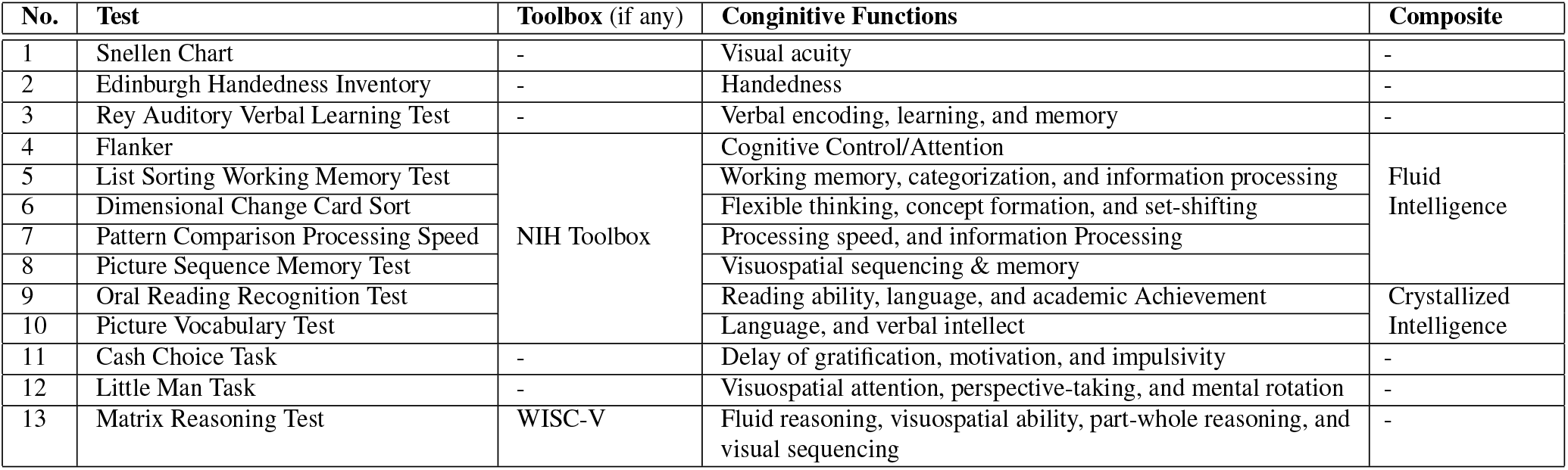
List of neurocognitive tests performed in ABCD initiative. Acronyms-WISC: Wechsler Intelligence Scale for Children, NIH: National Institute of Health

Brain development is influenced by both natural and nurture factors^5^. The former factor is coded in the gene. The latter factor is related to the environment, nutrition, socioeconomics, lifestyle, and other factors. Nature and nurture jointly shape the development of the human brain^6^, which can be observed noninvasively by magnetic resonance imaging (MRI)^7^. Variability in natural and nurtural factors causes individual differences in cognitive abilities, such as variation in the level of intelligence (i.e., measured in terms of the intelligence quotient (IQ)), attention, decision-making, memory, and executive function^4^. Several studies utilized ABCD data and showed the correlation between the neurocognition and parent-rated problem behaviors^4^, neurocognitive development and impacts of substance use^2^, the activation pattern of functional MRI and different neurocognitive processes (i.e., cognitive impulse control, reward anticipation and receipt, and working memory and emotion reactivity)^8^, suicide ideation and neurocognition^9^, and age and longitudinal stability of individual differences in neurocognition^10^. However, a comprehensive study to elucidate the effect of both natural and nurtural factors on adolescent neurocognition is yet to perform. These natural and nurtural factors are embedded in demographics and socio-economics, respectively, and also blended into the brain’s macro and microstructure. Thus, this article aimed to investigate how adolescent neurocognition is affected by demographic, socio-economic, and brain anatomical factors. Correlations between different neurocognitive test scores and variables were estimated using Pearson’s correlation. In addition, statistical significance is sought using the Bonferroni-adjusted^11^ *p*-values (i.e., *Q*-values) of 0.01.

## Results

### Experimental Setup

We used the Pearson correlation to estimate the correlation coefficient (*r* : [*−*1, 1]) between NIH toolbox neurocognitive subdomain test scores (see rows 4-10 of Table 1) and anatomical or demographic or socio-economic factors. The NIH toolbox also provides composite scores, namely fluid composite and crystallized composite, to allow for general evaluation of overall cognitive acuity. The fluid composite includes fluid ability measures such as Flanker, List Sorting Working Memory, Dimensional Change Cart Sort, Pattern Comparison Processing Speed, and Picture Sequence Memory (rows 4-8 of Table 1). On the other hand, the crystallized composite includes the Oral Reading Recognition and Picture Vocabulary Tests (rows 9-10 of Table 1). Fluid abilities are influenced by biological processes, which play an important role in adapting to novel situations in everyday life and are less dependent on past learning experiences^12^. On the contrary, crystallized abilities are known to be more dependent on past experience and less on biological influences^12^. Finally, the total composite score is estimated by averaging the fluid and crystallized composites. In this study, we also used Pearson’s correlation to estimate the correlation coefficient (*r* : [*−*1, 1]) between NIH toolbox neurocognitive composite test scores and anatomical or socio-economic factors. Furthermore, we used Bonferroni-adjusted *p*-values (i.e., *Q*-values) of 0.01 and plotted only those *r* values in Figs. 1–7 heatmaps that satified Q *≤* 0.01; otherwise left blank. In addition, we used several symbols in Figs. 1–7 heat maps with different colors to represent the correlation range in which a particular *r* falls.

**Figure 1.**
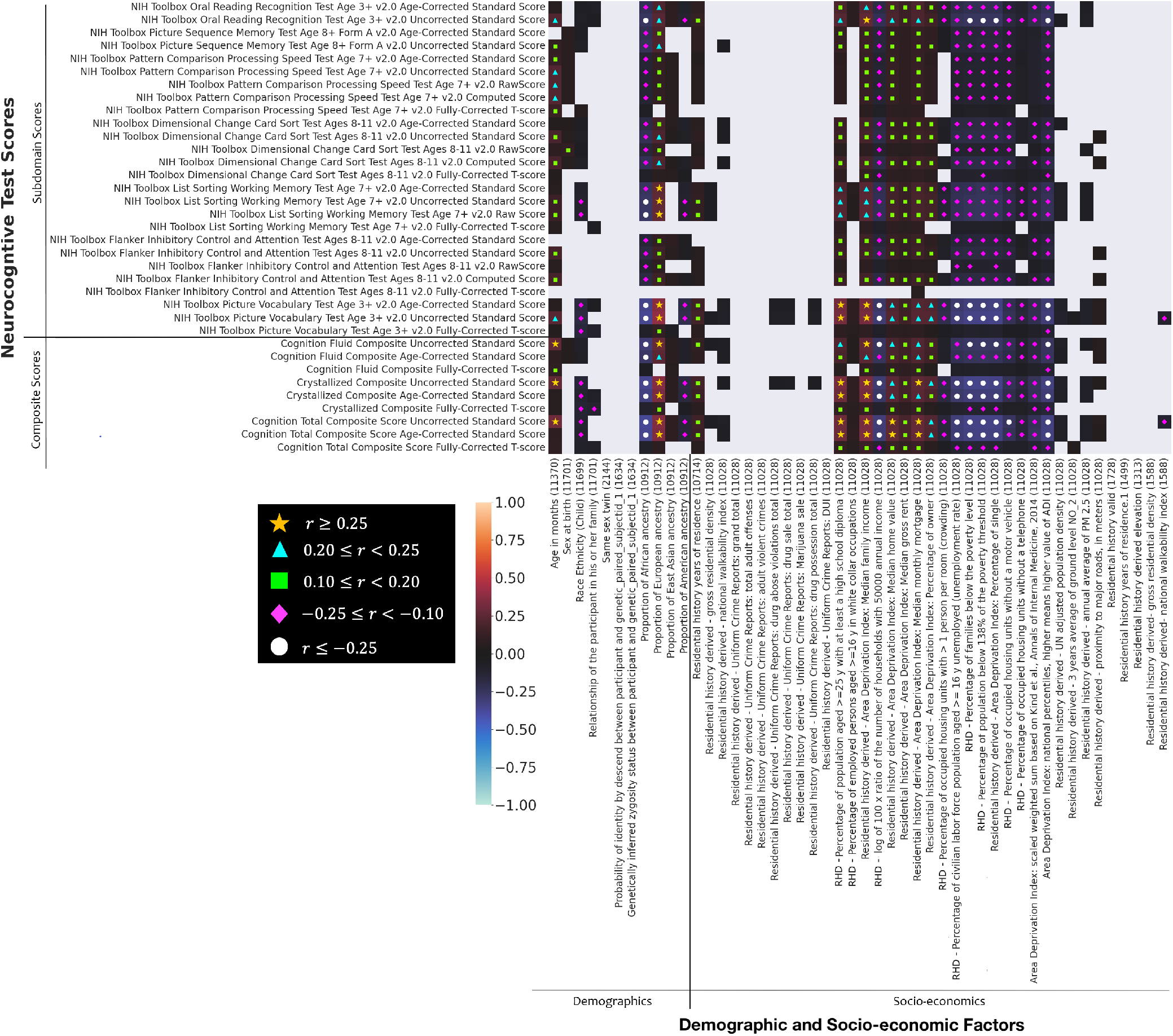
Heatmap showing the Pearson correlation coefficient (*r*) between neurocognitive test scores and demographic/ socio-economic factors. Statistically not significant correlation values (i.e., *Q >* 0.01) are left blank. Acronyms-RHD: residential history derived.

In the ABCD dataset, many subjects did not have a complete set of anatomical and socio-economic factor values on record. Therefore, we estimated the correlation between test scores and a factor for those subjects who have that particular factor in record and mentioned the number of subjects in parentheses on the x-axis. However, if any anatomical or socio-economic factor is available for less than 500 subjects, we did not include that factor in our correlation analysis. The ABCD dataset also comes with standard uncorrected scores, age-corrected standard scores, and fully corrected T scores from neurocognitive tests^12^. The uncorrected standard score represents the performance comparison between the test-taker (i.e., subject) and the entire NIH toolbox representative normative samples (normative mean = 100, standard deviation = 15) in the United States (US) population, regardless of any socio-economic factors. Age-corrected standard scores, on the other hand, compare the score of the test taker to those in the NIH toolbox nationally representative normative sample ‘at the same age,’ which were estimated separately for children (ages 3-17 years) and adults (ages 18-85 years). The fully corrected T-score (normative mean = 50, standard deviation = 10) compares the score of the test-taker to those in the NIH Toolbox nationally representative normative sample after the adjustment for age, gender, race/ethnicity, and educational attainment (for ages 3-17 years, parent’s level of education is used).

### Effect of Demographic and Socio-economic Factors on Neurocognition

In Fig. 1, we show the heat map of the Pearson correlation coefficient (*r*) between the scores of the neurocognitive test and the demographic/socio-economic factors. We see in this heat map that the uncorrected scores for fluid, crystal, and total composite are significantly correlated (*r >* 0.25) with age. One of the notable findings in this study is the effect of ancestry on different composite scores. We see that the proportion of African ancestry is significantly negatively correlated (*r < −*0.25) with different composite scores followed by the proportion of American ancestry (*−*0.25 *≤ r < −*0.10), while the proportion of

European ancestry is significantly positively correlated (*r >* 0.25) with different composite scores. Other factors that positively affect (*r >* 0.25) neurocognition acuity (as seen in terms of composite scores) are found to be the percentage of the population aged *≥* 25 years with high school diploma, median family income, median home value, and median monthly mortgage. In contrast, factors that negatively affect (*r >* 0.25) neurocognition acuity are found to be the percentage of unemployed civilian labor aged *≥* 16 years, the percentage of families below the poverty level, and the percentage of singles in a neighborhood. Other mildly negative factors (*−*0.25 *≤ r < −*0.10) affecting neurocognitive functions are the percentage of occupied housing units without a motor vehicle, the percentage of occupied housing units with *>* 1 person per room, and the percentage of occupied housing units without a telephone in a neighborhood.

### Effect of Cortical Areas of Brain ROIs on Neurocognition

In Fig. 2, we show the heat map of the Pearson correlation coefficient (*r*) between neurocognitive test scores and cortical areas (mm^2^) of brain region-of-interests (ROIs). We see in this heat map that the uncorrected and age-corrected crystallized composite and the total composite scores are positively correlated (0.10 *≤ r <* 0.20) with the cortical ROI areas that cover almost the entire brain. We also observe that the uncorrected and age-corrected subdomain tests scores such as the oral reading and the picture vocabulary test scores, which form the crystallized ability (see Table 1), are also positively correlated to (0.10 *≤ r <* 0.20) with the cortical ROI areas that cover almost the entire brain. However, fluid composite scores were not found to be strongly correlated with cortical ROI areas, although the working memory test scores show a positive correlation (0.10 *≤ r <* 0.20) with almost all cortical ROI areas in the brain. Furthermore, the total left, right, and whole brain cortical areas show a higher positive correlation (0.20 *≤ r <* 0.25) with the uncorrected and age-corrected crystallized composite and total composite scores.

**Figure 2.**
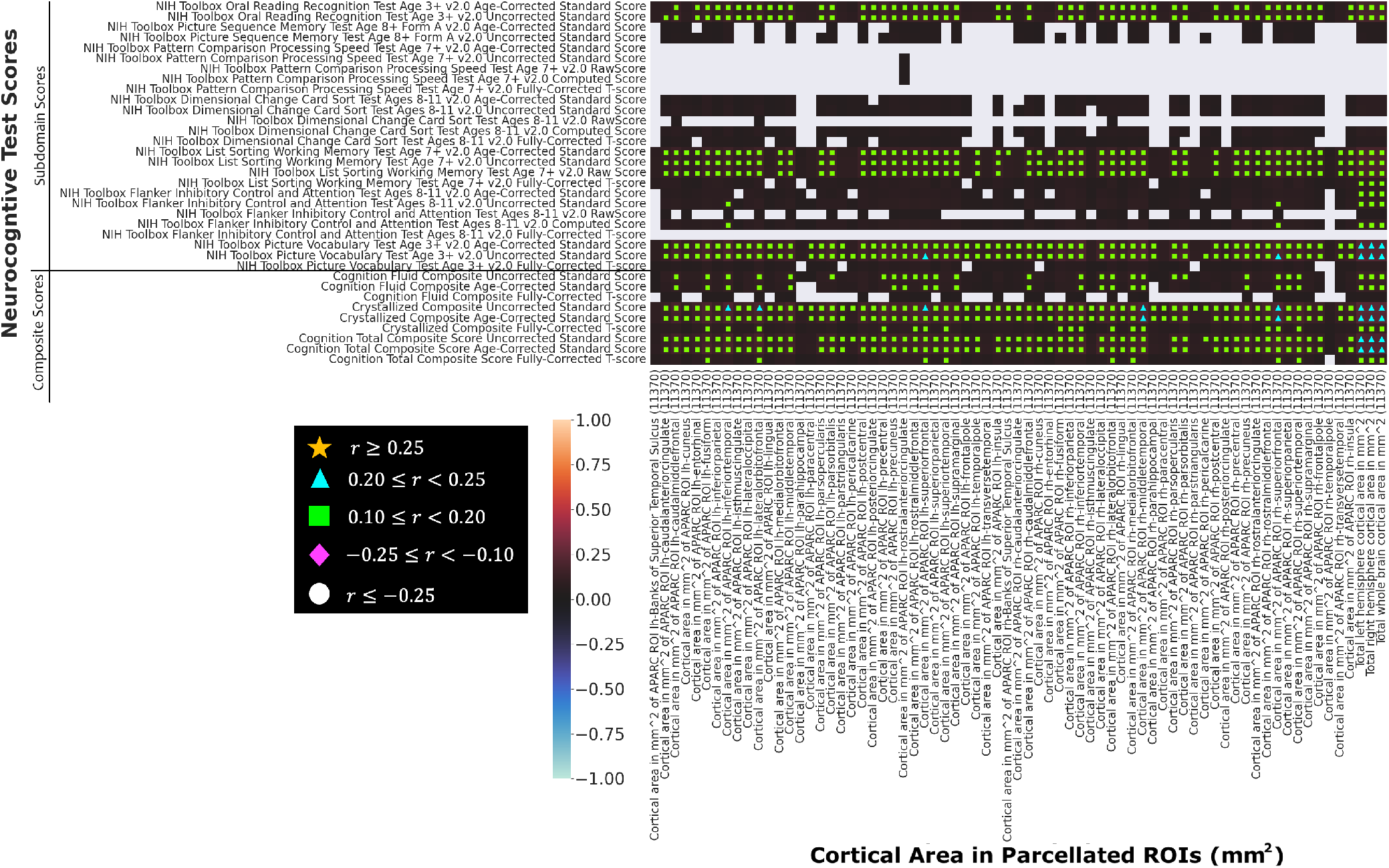
Heat map showing the Pearson correlation coefficient (*r*) between neurocogntive test scores and cortical areas (mm^2^) of brain ROIs. Statistically not significant correlation values (i.e., *Q >* 0.01) are left blank. Acronyms-APARC: automatic parcellation, lh: right-hemespheric, and rh: right-hemespheric.

### Effect of Cortical Thickness of Brain ROIs on Neurocognition

We show a heat map of the Pearson correlation coefficient (*r*) between neurocognitive test scores and cortical thickness (mm) of brain ROIs in Fig. 3. We see in this heat map that, unlike the cortical ROI areas, the cortical thickness in the brain ROIs does not show a strong trend of positive correlation with the neurocognitive test scores. However, the uncorrected and age-corrected crystallized composite and the total composite scores are positively correlated (0.10 *≤ r <* 0.20) with the cortical thickness in the left- and right-hemespheric lingual and parahippocampal ROIs, and right-hemespheric lateral occipital ROI.

**Figure 3.**
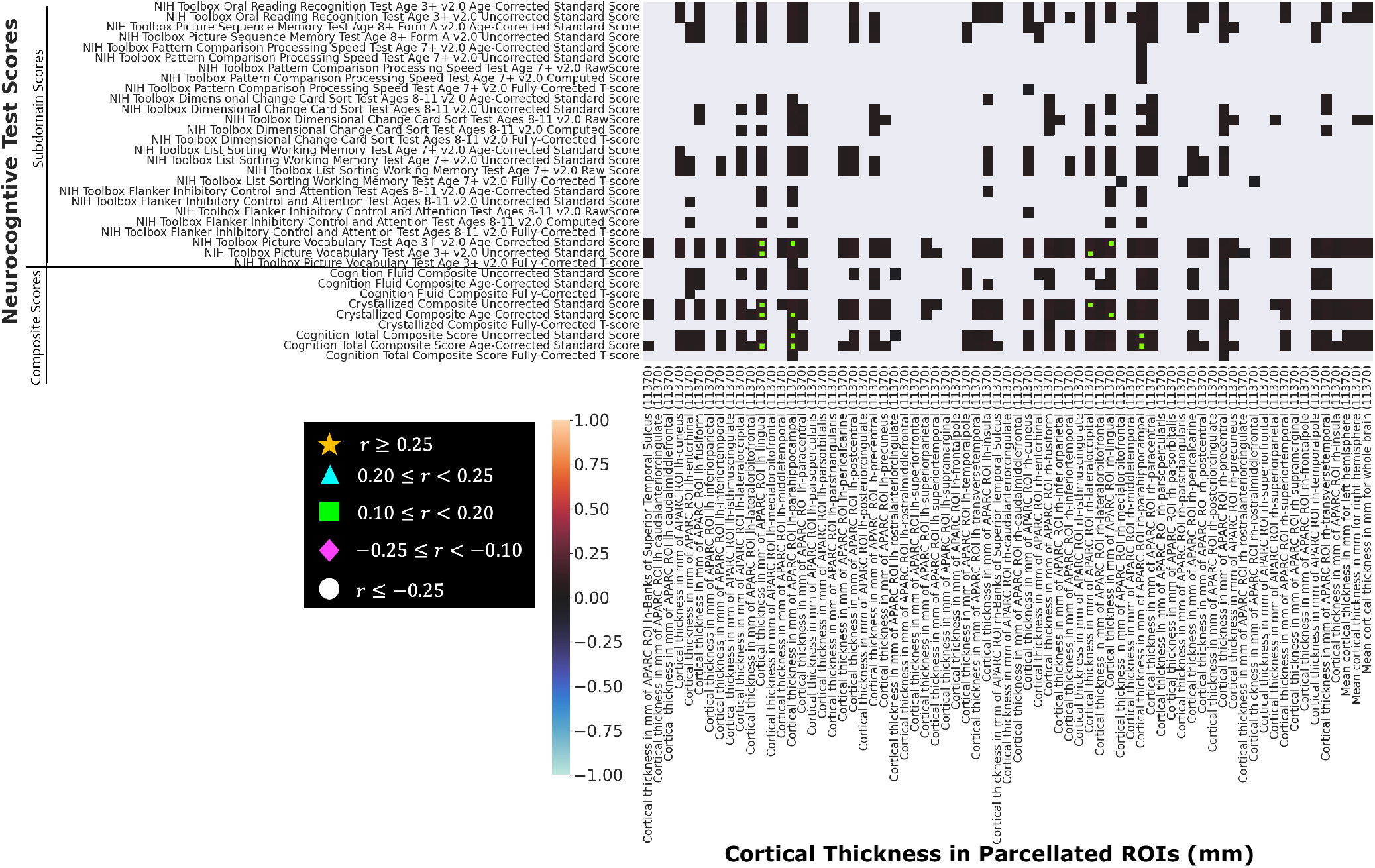
Heat map showing the Pearson correlation coefficient (*r*) between neurocognitive test scores and cortical thickness (mm) of brain ROIs. Statistically not significant correlation values (i.e., *Q >* 0.01) are left blank. Acronyms-APARC: automatic parcellation, lh: right-hemespheric, and rh: right-hemespheric.

### Effect of Cortical Volumes of Brain ROIs on Neurocognition

Furthermore, we show the heat map of the Pearson correlation coefficient (*r*) between neurocognitive test scores and cortical volumes (mm^3^) of brain ROIs in Fig. 4. Similar to cortical ROI areas, the cortical volumes in the brain ROIs show a strong trend of positive correlation with the uncorrected and age-corrected crystallized composite and total composite scores. We see in the heat map of Fig. 4 that the uncorrected and age-corrected crystallized composite and total composite scores are positively correlated (0.10 *≤ r <* 0.25) with the cortical ROI volumes that cover almost the entire brain. We also observe that the uncorrected and age-corrected subdomain tests scores such as the oral reading and the picture vocabulary test scores, which form the crystallized ability (see Table 1), are also positively correlated to (0.10 *≤ r <* 0.25) with the cortical ROI volumes that cover almost the entire brain. However, as with cortical ROI areas, fluid composite scores are not found to be strongly correlated with cortical ROI volumes, although list sorting working memory test scores show a positive correlation (0.10 *≤ r <* 0.25) with almost all cortical ROI areas of the brain. Furthermore, the total left, right, and whole brain cortical volumes show a higher positive correlation with the uncorrected and age-corrected crystallized composite (*r ≥* 0.25) and total composite scores (0.20 *≤ r <* 0.25).

**Figure 4.**
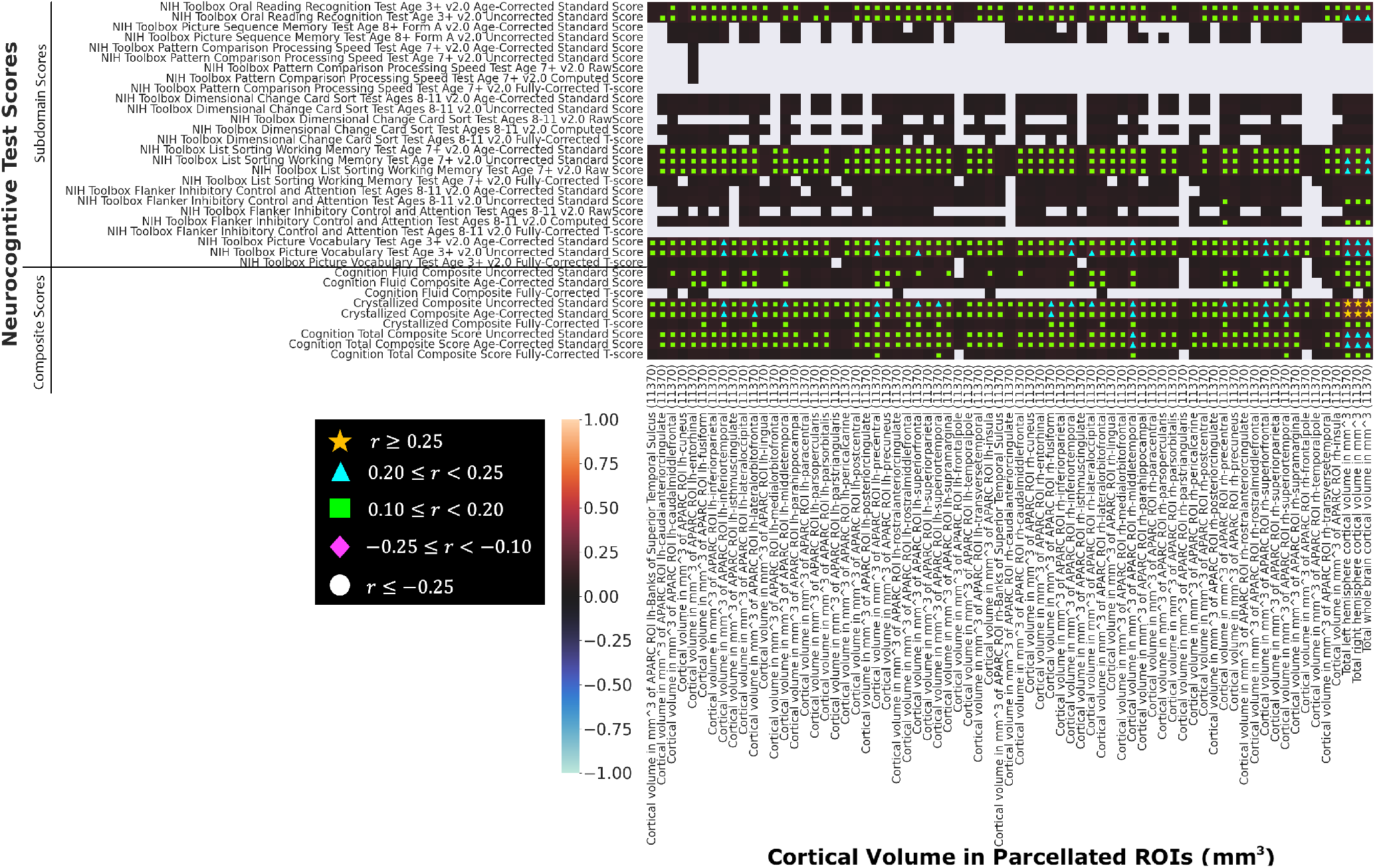
Heat map showing the Pearson correlation coefficient (*r*) between neurocogntive test scores and cortical volumes (mm^3^) of brain ROIs. Statistically not significant correlation values (i.e., *Q >* 0.01) are left blank. Acronyms-APARC: automatic parcellation, lh: right-hemespheric, and rh: right-hemespheric.

### Effect of Cortical Sulcal Depth of Brain ROIs on Neurocognition

We also show a heat map of the Pearson correlation coefficient (*r*) between neurocognitive test scores and cortical sulcal depth (mm) of brain ROIs in Fig. 5. We see in this figure that the uncorrected and age-corrected crystallized composite and total composite scores are positively correlated (0.10 *≤ r <* 0.20) with cortical sulcal depths in the left- and right-hemespheric medial orbito-frontal, left-hemespheric temporal pole, and righ-hemespheric superior frontal ROIs, while negatively correlated (*−*0.25 *≤ r < −*0.10) with cortical sulcal depths in the left- and right-hemespheric posterior cingulate, left- and right-hemespheric transverse temporal, and righ-hemespheric caudal anterior cingulate ROIs.

**Figure 5.**
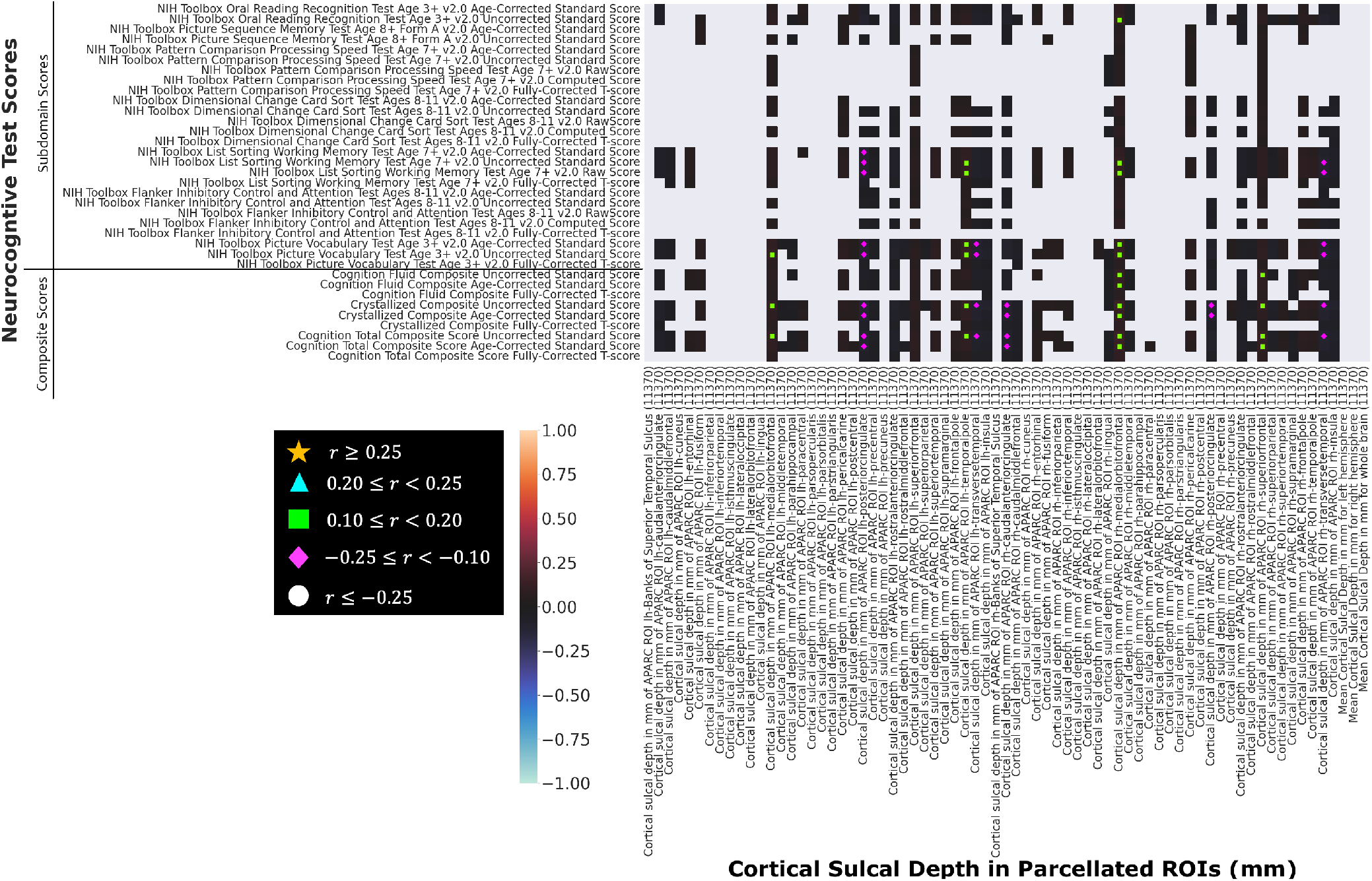
Heat map showing the Pearson correlation coefficient (*r*) between neurocognitive test scores and cortical sulcal depth (mm) of brain ROIs. Statistically not significant correlation values (i.e., *Q >* 0.01) are left blank. Acronyms-APARC: automatic parcellation, lh: right-hemespheric, and rh: right-hemespheric.

### Effect of Subcortical Volumes of Brain ROIs on Neurocognition

Additionally, we show the heat map of the Pearson correlation coefficient (*r*) between neurocognitive test scores and subcortical volumes (mm^3^) of brain ROIs in Fig. 6. Similar to cortical ROI areas and volumes, the subcortical volumes in the brain ROIs show a strong trend of positive correlation with the uncorrected and age-corrected crystallized composite and total composite scores. We see in the heat map of Fig. 6 that the uncorrected and age-corrected crystallized composite and total composite scores are positively correlated (*r ≥* 0.10) with the subcortical ROI volumes that cover almost the entire brain. We also observe that the uncorrected and age-corrected subdomain tests scores such as the oral reading and the picture vocabulary test scores, which form the crystallized ability (see Table 1), are also positively correlated to (*r ≥* 0.10) with the subcortical ROI volumes that cover almost the entire brain. However, fluid composite scores are positively correlated with subcortical volumes in sparsely located ROIs, although the list sorting working memory test scores show a positive correlation (0.10 *≤ r <* 0.20) with almost all subcortical brain ROI volumes. Furthermore, subcortical volumes in the entire brain, supratentorial, and subcortical gray areas show a higher positive correlation with the uncorrected crystallized composite score (*r ≥* 0.25).

**Figure 6.**
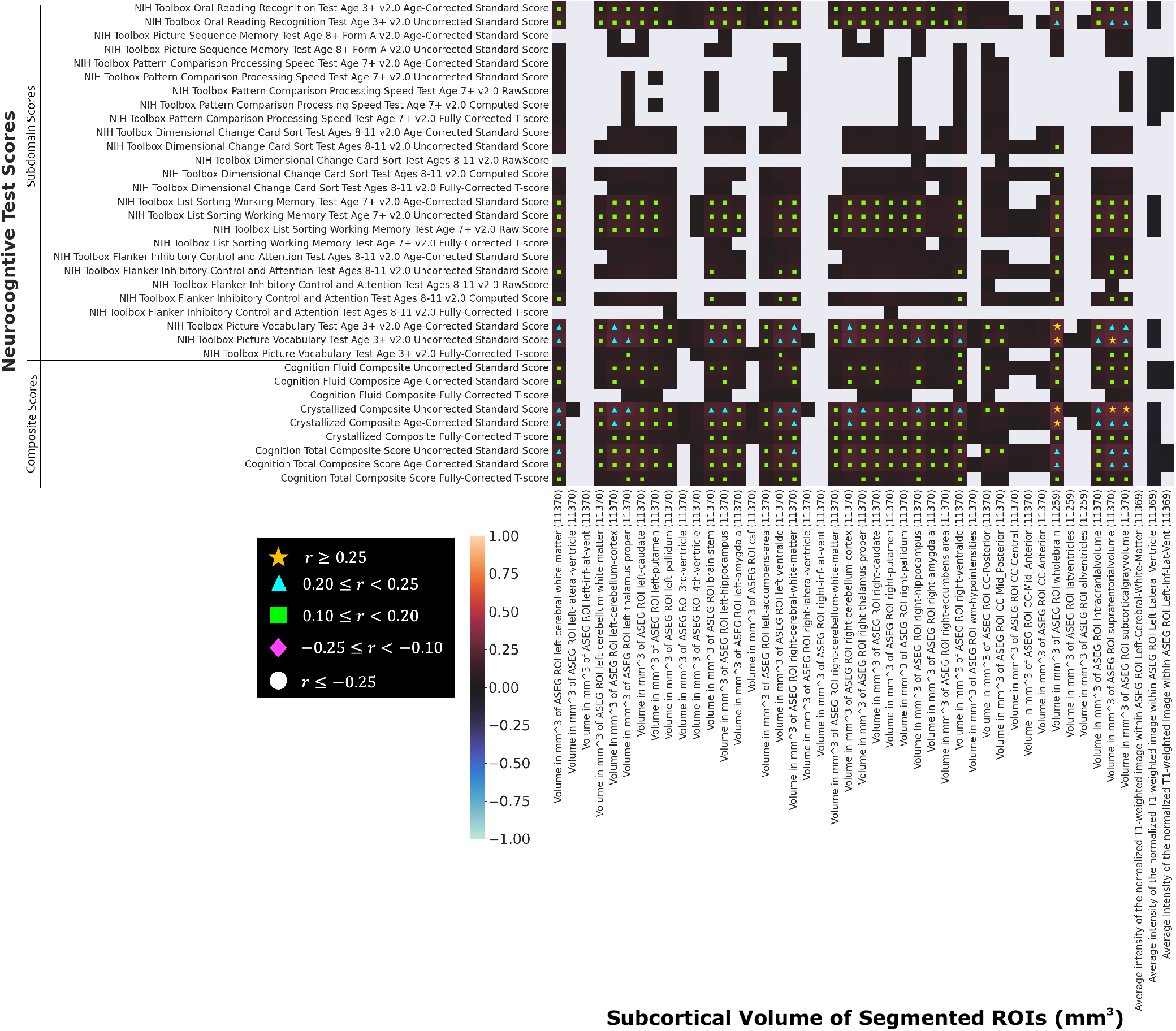
Heat map showing the Pearson correlation coefficient (*r*) between neurocogntive test scores and subcortical volumes (mm^3^) of brain ROIs. Statistically not significant correlation values (i.e., *Q >* 0.01) are left blank. Acronyms-ASEG: automatic segmentation.

### Effect of Average T1- and T2-weighted Intensity of Brain ROIs on Neurocognition

Finally, we show the heat map of the Pearson correlation coefficient (*r*) between neurocognitive test scores and average T1- and T2-weighted intensity of brain ROIs in Fig. 7. This figure shows that the average intensities of brain ROIs do not strongly contribute to neurocognitive acuity. The only notable correlation is observed between the volume of the right pallidum and the age-corrected pattern comparison processing speed with *−*0.25 *≤ r < −*0.10.

**Figure 7.**
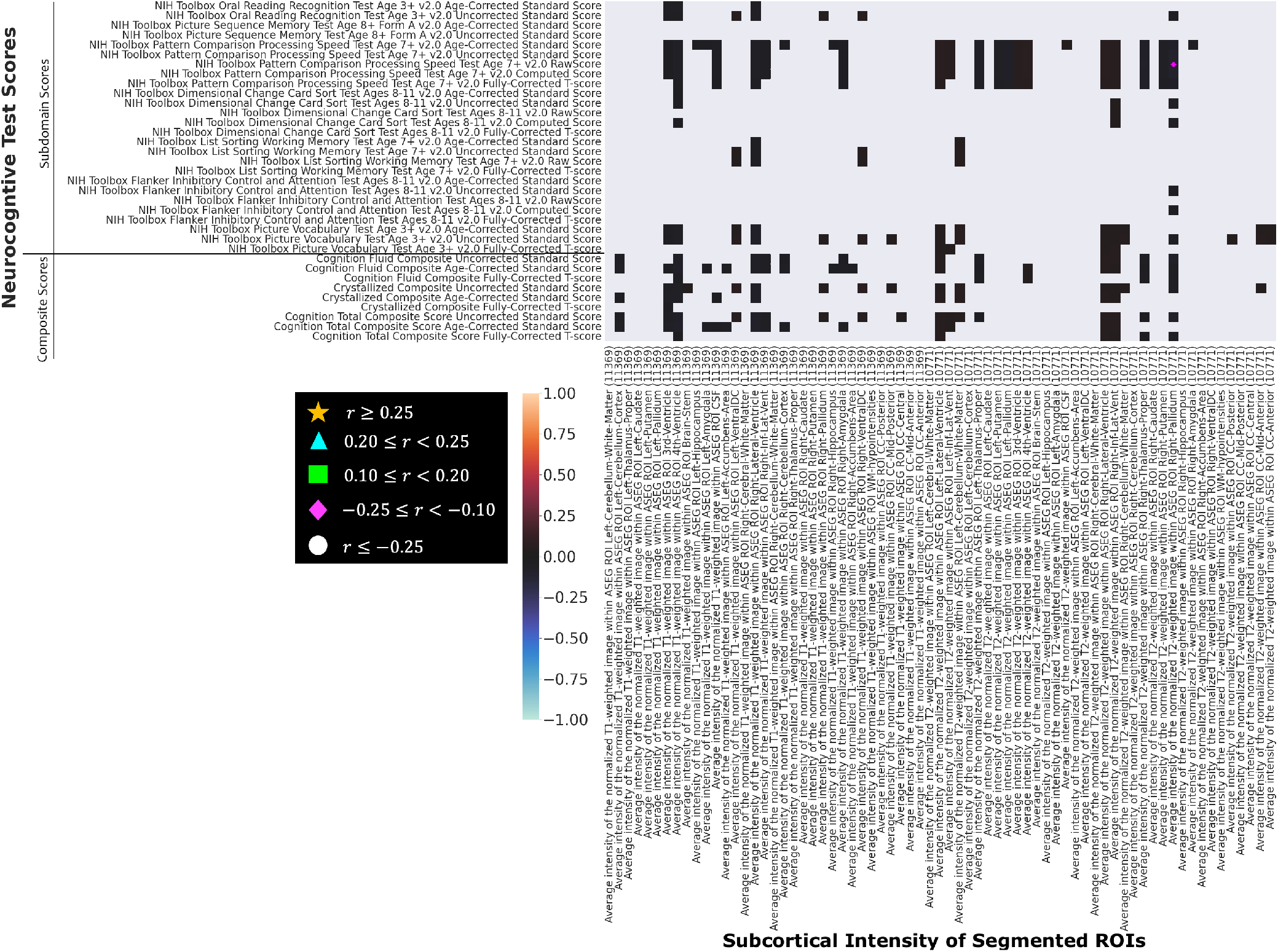
Heat map showing the Pearson correlation coefficient (*r*) between neurocogntive test scores and average T1- and T2-weighted intensity of brain ROIs. Statistically not significant correlation values (i.e., *Q >* 0.01) are left blank. Acronyms-ASEG: automatic segmentation.

## Discussion

This Pearson correlation-based study analyzed the effect of brain structure, demographics, and socio-economics on adolescents’ neurocognitive functions and revealed several key findings. At the same time, the analysis reveals some open research questions. One of the major observations in this study is the interrelation of neurocognitive functions with the ancestry of adolescents, as seen in Fig. 1. We observed a strong positive correlation (*r ≥* 0.25; *Q ≤* 0.01) of European ancestry with uncorrected fluid, crystallized, and total composite scores. In contrast, we observed a strong negative correlation (*r ≤ −*0.25; *Q ≤* 0.01) of African ancestry with uncorrected fluid, crystallized, and total composite scores. In addition, we observed a moderate negative correlation (*−*0.25 *≤ r < −*0.10; *Q ≤* 0.01) of American ancestry and almost no correlation of East Asian ancestry with the uncorrected crystallized and total composite scores (see Fig. 1). However, one of the remaining questions is *how a mixed ancestry (i*.*e*., *father and mother are of different ancestral lineage) would contribute to different neurocognitive subdomains and composite abilities*.

We also observed in Fig. 1 for the sample size of 11,701 that sex does not affect any neurocognitive functions in adolescents, except executive function (Dimensional Change Card Sort Test; 0.10 *≤ r <* 0.20; *Q ≤* 0.01). Similar findings are reported by French et al.^13^ on a sample size of 2,845 (10-18 years). However, another study^14^ on a sample size of 3,500 (8-21 years) showed that sex difference has a smaller but noticeable effect on all social cognition tests. Furthermore, sex is an important determinant of neurocognitive impairments in diseased populations; for example, populations with the human immunodeficiency virus (HIV)^15^. Roalf et al.^3^ studied 9,138 youths (8-21 years) and reported that within-individual variability of neurocognitive functions is the highest in childhood, declines into mid-adolescence, and increases again into adulthood. This finding agrees with our findings. Therefore, another open question would be *why the effect of sex on neurocognitive functions diminishes for a short span of time during adolescence (age ∼8–18 years)*.

In Fig. 1, we further observed that the disadvantage of the neighborhood in terms of crime reports for adult offenses, violent crimes, drug abuse, and drug sales does not affect any neurocognitive functions in adolescents aged 9 to 10 years. However, the disadvantage of the neighborhood in terms of the percentage of the population (aged *≥* 16 years) unemployed, the percentage of families below the poverty level, and the percentage of the population being single adversely affects (*r ≤ −*0.10; *Q ≤* 0.01) neurocognitive functions in adolescents. In contrast, the advantage of the neighborhood in terms of the percentage of the population (aged *≥* 25 years) having at least a high school diploma, median family income, median home value, and median monthly mortgage favorably affects (*r ≥* 0.10; *Q ≤* 0.01) the neurocognitive functions in adolescents (see Fig. 1). In this context, it would be important to find out using multivariate correlation analysis *how neurocognitive functions in adolescents are affected when they live in a mix of neighborhood advantages and disadvantages factors*.

In addition to the effects of demographic and socio-economic factors, brain structure has shown considerable effects on neurocognitive functions in adolescents. However, the degree of effects by different brain regions and geometry varies. In Figs. 2, 3, 4, 5, we showed heat maps of the Pearson correlation coefficient (*r*) between neurocognitive test scores and cortical areas, cortical thickness, cortical volumes, and cortical sulcal depths, respectively, for the same set of parcellated cortical gray matter ROIs. For cortical ROI areas, we see in Fig. 2 that the crystallized composite and the total composite scores are positively correlated (*r ≥* 0.10; *Q ≤* 0.01) with ROI areas that cover almost the entire brain. However, the cortical thickness for the same set of ROIs did not show a strong trend of positive correlation with the neurocognitive test scores (see Fig. 3). The cortical ROI volume (product of cortical area and thickness) for the same set of ROIs, on the other hand, showed a similar trend of positive correlation (*r ≥* 0.10; *Q ≤* 0.01) with neurocognitive test scores as cortical areas (see Figs. 2 and 4). In contrast, cortical sulcal depth for a few sparsely located ROIs showed a moderate positive (0.10 *≤ r <* 0.20; *Q ≤* 0.01) and negative (*−*0.25 *≤ r < −*0.10; *Q ≤* 0.01) correlation with the neurocognitive test scores (see Fig. 5). Similarly to the cortical anatomical structure, the subcortical structure plays an important role in neurocognitive development in adolescents. We see in Fig. 6 that the crystallized composite and the total composite scores are positively correlated (*r ≥* 0.10; *Q ≤* 0.01) with subcortical ROI volumes that cover almost the entire brain. However, MRI contrast of the subcortical tissues did not show any significance for neurocognitive development in adolescents, as evidenced by the poor correlation between neurocognitive test scores and the average intensity of T1 and T2-MRI of subcortical ROIs in Fig. 7. Our overall observation on Figs. 2-7 depicts that the anatomical structure of brain tissue plays a more significant role in crystallized cognition than fluid cognition ability. On the other hand, the brain network homogeneity correlates with fluid intelligence in children^16^. Therefore, it warrants investigating *whether the role of functional brain connectivity and/or biochemical diffusivity in the development of fluid cognition is stronger than cortical and subcortical structures or similar*.

Our study also shed some light on the effects of natural *vs*. nurtural agents on the development of neurocognition in adolescents. Considering demographic and socio-economic factors in Fig. 1 as natural and nurtural factors, respectively, we see a clear comparative picture of the effects of these types of factors on adolescent neurocognition. However, anatomical and microstructural development in the brain is known to be governed by both natural (i.e., intrinsic molecular cues derived from gene expression) and nurtural (i.e., extrinsic input from sources outside of the organism) factors^17^. Thus, it would be interesting to investigate *what degree of change in brain structure does nature and nurture cause separately, and how it affects adolescent neurocognition*.

Last but not least, our correlation analysis was univariate, where demographic, socio-economic, and brain structural factors were individually used as a variable and correlated with different neurocognitive test scores. However, to analyze the covariance of multiple factors in the development of neurocognition in adolescents, it is necessary to implement *multivariate correlation analysis*. Furthermore, it is also necessary to investigate *the comparative roles of individual factors in terms of feature importance*, when used in a multivariate analysis.

## Methods

### Data

We accessed demographic, socio-economic, structural brain anatomy, and neurocognitive test scores from 11,878 samples from the Adolescent Brain Cognitive Development (ABCD) initiative^1^. We summarize the demographics of all subjects included in this study in Table 2. These data are collected in 21 sites across the United States using either of Siemens (Prisma or Prisma Fit), General Electric (MR 750), or Philips (Achieva dStream or Ingenia) MRI scanners with 3T magnetization. The MRI preprocessing steps^18^ involved (i) correction of gradient nonlinearity distortions in T1 and T2-MR images using scanner-specific nonlinear transformations, (ii) registration of T2-MR images to T1-MR images using mutual information, (iii) correction of intensity inhomogeneity correction B1-bias fields, (iv) rigid registration and resampling into alignment of MR images to an averaged reference brain in standard space, (v) cortical surface reconstruction and subcortical segmentation using FreeSurfer v5.3, and (vi) atlas-based automatic volumetric segmentation/parcellation and labeling of cortical and subcortical structures.

**Table 2.** Demographics for all participants included in this study for correlation analysis.

| Measure | Category | Number of Samples |
| --- | --- | --- |
| Total Samples | - | $N = 11,878$ (100%) |
| Age (years)<br>(mean: $9.91 \pm 0.62$) | 9 years | 6,077 (51%) |
|  | 10 years | 5,457 (46%) |
|  | Not available | 344 (3%) |
| Sex at birth | Male | 6,196 (52%) |
|  | Female | 5,679 (48%) |
|  | Other | 3 (Negligible) |
| Race/ethnicity | White | 6,180 (52%) |
|  | African American | 1,784 (15%) |
|  | Hispanic | 2,410 (20%) |
|  | Asian | 252 (2%) |
|  | Other | 1,247 (11%) |
|  | Not available | 5 (Negligible) |

### Pearson Correlation

The Pearson correlation measures the linear relationship between the attributes of two datasets *A* and *B* and produces a coefficient (*r*) value in the range of [*−*1, 1]. A value of *r* = *−*1 represents a perfect negative correlation, a value of *r* = +1 represents a perfect positive correlation, and a value of *r* = 0 represents no correlation between *A* and *B*. Although *r* = 0 indicates that there is no linear relationship between *A* and *B*, there may still be a higher-order relationship between the same datasets. The Pearson correlation between paired datasets (*A, B*) : *{*(*a*_1_, *b*_1_), (*a*_2_, *b*_2_), …, (*a*_*N*_, *b*_*N*_)*}* is mathematically defined as^19^:

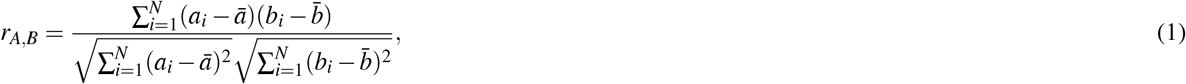

where 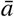 and 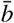 are the mean of all data points in datasets *A* and *B*, respectively.

